# Dairy manure analysis reveals significant risk of Antibiotic resistance from Extracellular DNA in Manure storage Pit

**DOI:** 10.1101/2025.11.07.687174

**Authors:** Najmuj Sakib, Daniel Andersen, Laura Jarboe, Adina Howe

**Author notes:** Corresponding Author: Adina Howe, Department of Agricultural and Biosystems Engineering, Iowa State University, Ames, IA 50011, United States.

## Abstract

Dairy manure pit storage systems are significant reservoirs for antimicrobial resistance genes (ARGs). These genes occur in both intracellular DNA (iDNA) and extracellular DNA (exDNA), but their distribution across these categories in fresh (loafing pen surface) and pit-stored dairy manure has not been previously characterized. To address this gap, we quantified the abundance of six ARGs (*tetG*, *tetM*, *tetX-* tetracyline, *sul1*-sulfonamide, and *ermB-*macrolide) and three mobile genetic elements (MGEs) (*intI1*, *intI2*, and *intI3*) in iDNA and exDNA extracted from fresh and pit-stored manure collected at a dairy farm in Iowa. While total DNA yields were lower in pit-stored relative to fresh manure samples, exDNA-to-iDNA ratios were significantly elevated across all genes (p<0.001), indicating relative enrichment of exDNA during storage. Notably, *tetM* exhibited a higher free (unattached) to bound (surface-attached) exDNA ratio in pit samples, which suggested an increased potential for gene transfer in the pit. Correlation network analysis revealed similar numbers of strong ARG–MGE associations in pit and fresh exDNA, but lower interconnectivity in pit exDNA. Merging fresh and pit datasets showed wider ARG-MGE associations: *intI1* and *intI3* strongly co-occurred with tetracyclines and macrolide resistance in iDNA, while *sul1* correlated with MGEs only in the exDNA network. Microbial community profiling showed similar taxa in exDNA across manure types, while iDNA communities diverged significantly. This result could support that exDNA is relatively stable over time and in varying environments, and that iDNA is relatively more reflective of selective pressures. Overall, our results highlight exDNA as a critical but overlooked reservoir of resistance determinants, warranting further investigation and targeted management strategies in dairy systems.

**Importance:** To date, extracellular DNA (exDNA) has been shown to contribute to the spread of antibiotic resistance genes (ARGs) in the environment; however, few studies have evaluated its enrichment in dairy pit-stored manure systems. This study demonstrates that dairy manure pits concentrate exDNA during dairy manure storage and serve as a reservoir for ARGs, along with mobile genetic elements that can facilitate subsequent gene transfer. The results of this study are a strong rationale for further investigation and targeted management strategies of exDNA in manure pits.

## Introduction

The use of antibiotics in livestock is crucial for animal health but has been associated with increased selective pressure for antibiotic resistance and the enrichment of antibiotic-resistant bacteria (ARB) and genes (ARG) in animal guts and manures (Winckler and Grafe, 2001; Looft et al., 2012). The World Health Organization has reported that up to 80% of total antibiotic consumption occurs in the animal sector (WHO, 2017), and this demand is projected to grow globally, as antimicrobial sales are expected to rise 11.5% by 2030 across all continents (Tiseo et al., 2020). In the United States’ dairy industry, it is estimated that up to 90% of operations use intramammary antibiotics, especially β-lactams (penicillins and cephalosporins) and tetracyclines, primarily for dry cow therapy (Oliver et al., 2020). While antibiotic use in livestock is essential for animal health, its widespread application creates downstream challenges beyond the animal gut. One critical point of concern is manure management, where slurry manure pits have been identified as hotspots for antimicrobial resistance (Lima et al., 2020). Pit storage systems are often favored in regions with high nutrient demand for crop growth because they help maintain higher nutrient levels, making them integral to nutrient recycling strategies (Worley, 2015). However, these storage pits accumulate fresh manure rich in ARB and ARGs, along with wastewater inputs that introduce additional chemical stressors. This wastewater can contain chemicals, such as heavy metals (like copper and zinc from footbaths), or other antimicrobials, that can also impact the development of antimicrobial resistance in manure pit microbial communities (Todman et al., 2024).

Within dairy manure as well as animal manures broadly, many studies have focused on characterizing AMR risks through the identification and quantification of ARGs in manure and manure pits (Wichmann et al., 2014; Lima et al., 2020; Oliver et al., 2020; Baker et al., 2022). The emphasis of these studies has been on intracellular DNA (iDNA) encoding for ARGs. More recently, extracellular DNA (exDNA) has been identified as playing an increased role in the dissemination of antibiotic resistance (Sui et al., 2019; Zarei-Baygi and Smith, 2021; Wang et al., 2024). ExDNA originates from the lysis of dead bacteria or is actively secreted by living cells. Within a microbial community, ARGs can be contained in iDNA, exDNA, or both. Thus, there can be both cell-associated intracellular ARGs (iARGs) and extracellular ARGs (exARGs). ExDNA is generally considered free and readily accessible to competent cells (Nagler et al., 2018b; Yuan et al., 2019), and there are concerns that exARGs may be taken up by competent bacterial cells and genes that encode for antibiotic resistance between bacteria (Woegerbauer et al., 2020).

Beyond their inherent accessibility, the persistence and mobility of exARGs can be influenced by environmental and operational factors within manure systems. For example, dairy cleaning agents such as disinfectants can accelerate exDNA release and transformation (Zhang et al., 2017; Jin et al., 2020), while mobile genetic elements (MGEs) like integrons further facilitate gene transfer (Gillings et al., 2015). This suggests that manure pits containing abundant exARGs and MGEs could be hotspots for resistance dissemination. While studies on exARGs in various environments are available, they primarily focused on sludge and compost from livestock (Zhang et al., 2013; Zou et al., 2022; Tang et al., 2023; Xu et al., 2023; Yu et al., 2023; Bao et al., 2024; Xin et al., 2025), wastewater (Sui et al., 2019; Xin et al., 2024), manure-amended soil (McKinney et al., 2018; McKinney and Dungan, 2020), and fertilizers (Goetsch et al., 2020; Liu et al., 2024). Manure storage pits containing abundant exARGs and MGEs could also serve as hotspots for resistance dissemination, however the extent to which these factors influence gene transfer remains poorly understood.

In this study, we evaluate the hypotheses that the chemical and biological pressures of manure pits result in a greater enrichment of ARGs compared to fresh manure and that this enrichment is greater in exARGs compared to iARGs. To test this hypothesis, samples were compared between fresh manure and the manure storage pit of an operational dairy farm. Within each sample, ARGs were classified as iARGs or exARGs based on their location within or external to bacterial cells. In exDNA fractions, we further estimated the quantities of free versus bound fractions of exARGs. We particularly focused on characterizing selected ARGs previously identified in dairy manures, genes associated with resistance to macrolides (*ermB*), sulfonamides (*sul1*), and tetracyclines (*tet33, tetG, tetM, tetX*) (He et al., 2020). We also identified mobile genetic elements (MGEs) that may influence ARG transfer, particularly integrons (*intI1, intI2, intI3*). Finally, we compared the microbial communities represented in both iDNA and exDNA fractions from the same samples to better understand the origin of these ARGs. The outputs of this study are the first to provide a characterization of exARGs and iARGs in dairy pit-stored manure and comparing their AMR risks relative to fresh manure.

## Methods

### Site description, sample collection and processing

Fresh manure was collected in July 2023 from the loafing floor of a dairy (lactating cows) barn located at the Iowa State University Dairy Research Farm (GPS coordinates: 41.9777061, - 93.6494022). These cows have previously been treated with antibiotics, though no antibiotic were applied to feed. There were 420 milking cows at the time of sampling; among those, 90.3 % were Holstein, and 6-8% were Jersey. Manure was collected as a slurry sample from a reception pit prior to solid-liquid separation; the pit is located approximately 20 meters from the barn and receives manure (short-term storage of <6 hours) from pen surfaces and parlor and milkhouse washwater. For DNA extraction, three biological samples of fresh and pit manure were obtained. Fresh manure was a composite of 3-5 subsamples from different pen areas and contained minimal straw bedding. The solids content for fresh manure and slurry was 12 wt% and 4%, respectively. Samples were then placed in a 500 ml screw-cap plastic container, and it was filled up to three-fourths full to accommodate any gas creation. A portion of the samples was immediately processed for DNA extraction, and the rest was divided to be kept at 4°C for short-term usage and at –80°C for long-term storage.

### iDNA and exDNA extraction

To separate iDNA and exDNA, the previously described fractionation method (Nagler et al., 2018b) was used, resulting in the following fractions of exDNA: free (i.e., not attached to any surfaces), weakly bound (i.e., lightly attached to surfaces), and tightly bound (i.e., tightly attached to surfaces). After fractionation, DNA was extracted using the DNeasy PowerSoil Pro kit (Qiagen Laboratories, Germantown, MD, USA) (**Supplementary method M1**). To estimate the extraction efficiency for extracellular DNA, a subset of samples were spiked with live whole cells following the protocol of Mckinney et al., 2020 (McKinney and Dungan, 2020). Briefly, sub-samples of 100 mg (dry weight equivalent) manure (fresh or pit) were spiked with 100 ul whole cells (∼10^8^ cfu/ml, OD_600_ =1.8, 17.5 hours overnight culture) of a *Escherichia coli* (*E. coli* MG1655) encoding green fluorescence protein gene (*gfp*). (obtained from Ichiro Matsumura, Addgene plasmid # 26702 ; http://n2t.net/addgene:26702). DNA was extracted with the same methods as described above. Plasmids were extracted using Quick plasmid Miniprep Kit (Thermo Fisher Scientific, Waltham, MA, USA) and transformed into *E. coli* via electroporation. Quantitative real-time PCR (qPCR) was used to quantify the *gfp* copies in the extracted exDNA. A separate DNA extraction of transformed *E. coli* cells was carried out to quantify the initial *gfp* copies per ml of the sample. Extraction efficiency was then calculated as the ratio of *gfp* copies recovered from spiked manure samples to the number of *gfp* copies originally spiked into the samples. Unlike the protocol of Mckinney et al., 2020 which included DNA spiking, only whole-cell spiking was done for the current study. The DNA extraction yield and quality were verified with spectrophotometry (Nanodrop 2000 and Quant-it dsDNA Assay kit, High sensitivity, Thermo Fisher Scientific) and gel electrophoresis.

### Quantitative real-time qPCR for MGEs and ARGs

All extracted DNA samples were diluted to 1-2 ng/ul so that all measured concentrations fell within the range of known standards. qPCR assays were performed using previously described primers targeting *intI1*, *intI2*, *intI3*, *ermB*, *sul1*, *tet33*, *tetG*, *tetM* and *tetX* genes (Stedtfeld et al., 2018). The reactions were performed using a 96-well plate on a CFX96 Touch Real-time qPCR detection system (Biorad) using SYBR® Green approach. Templates of DNA standards were synthesized using gBlocks Gene fragments (IDT). Detailed information on the primers, templates and thermocyling conditions are listed in **Supplementary Tables S1**, **S2** and **S3**.

Standard templates were diluted to yield a series of seven 10-fold concentrations and subsequently used for qPCR standard curves (Ritalahti et al., 2006). Each gene was quantified in triplicate as well as with a standard curve and negative control. The limit of quantification (LOQ) was defined as the lowest standard concentration (the most diluted) of the linear range of the standard curve (Laht et al., 2014). In the rare instances where quantifications were observed in the negative controls, their Ct values were at least 3.3 cycles higher than the LOQ Ct values, ensuring they were well above the quantifiable range (**Supplementary Table S4**). All MGE and ARG qPCR results were normalized per gram (g) of dry manure (i.e., absolute abundance). Then, moisture contents were determined gravimetrically (McKinney et al., 2018) to normalize the gene data per gram (g) of dry manure for both sample types (details in **Supplementary Method M2**).

### 16s rRNA amplicon sequencing

16S rRNA amplicon sequencing for both types of DNA was performed by Argonne National Laboratory, utilizing an Illumina MiSeq platform with primers 515F and 806R. The sequencing run configuration was 151bp x 12bp x 151bp, including adapter sequences for Illumina flowcell (Caporaso et al., 2011, 2012; Walters et al., 2015). Further details are available in **Supplementary Method M3**.

The raw sequences were first processed for data analysis using DADA2 v1.16 to generate high-resolution amplicon sequence variants (ASVs)(Callahan et al., 2016). Taxonomic classification of these ASVs was performed by comparing ASV sequences against the SILVA (v138) database (McLaren and Callahan, 2021). The resulting data were further analyzed in R using the Phyloseq package. Alpha diversity was quantified using the Shannon index through the vegan package (Oksanen et al., 2024), with differences between DNA types assessed via Wilcoxon rank-sum tests. To quantify the magnitude of differences between groups, the effect size was calculated using Cliff’s delta when necessary. For beta diversity, Bray-Curtis distances were calculated with vegan (Oksanen et al., 2024) and evaluated by using permutational multivariate ANOVA with 999 permutations.

### Data analysis and statistics

All analyses were performed using R software (version 4.3.3). Graphs were generated using the ggplot2 package. All data were log-transformed and checked for normality (Shapiro-Wilk test) and equal variance (Levene’s test) to select the appropriate statistical tests for analysis. The gene copy values for free, weakly bound, and tightly bound DNA were combined and averaged to quantify the total extracellular DNA for each sample. For pairwise comparisons, Welch’s t-test was employed when data met the normality assumption; otherwise, the non-parametric Wilcoxon rank-sum test was used. To evaluate differences in gene copy numbers, relative abundances, or ratios across different antibiotic resistance genes (ARGs) within fresh manure or pit manure samples, one-way ANOVA was conducted for normally distributed data, while the Kruskal-Wallis test was applied as a non-parametric alternative. In these analyses, gene copy numbers, relative abundances, or ratios served as dependent variables. For all statistical tests, differences were considered significant at P < 0.05. Kendall’s Tau correlation test was also performed to assess the relationship between the absolute abundances of ARGs and MGEs. This non-parametric measure was chosen to account for potential non-linear relationships and to handle tied ranks in the data.

## Results

### Manure storage pit depletes iDNA relative to exDNA

Fresh manure and manure pit samples were taken from an active dairy farm, and iDNA and exDNA were extracted. The extracted iDNA yields ranged from 23.9–68.3µg/g of sample dry weight, whereas exDNA yields-derived from free, weakly bound and tightly bound fractions-were lower (0.8–10.4 µg/g). Both iDNA and exDNA were less abundant in pit samples compared to fresh manure (**Table 1**). To validate the exDNA extraction method, known quantities of whole cell *E. coli* carrying the *gfp* gene were spiked into samples. A low recovery of the *gfp* gene in the exDNA pool (0.003%–0.02%, *P* < 0.05, one-sample t-test) indicated that the extraction method effectively minimizes cell lysis and extraction of intracellular DNA.

**Table 1.** Extracted yields (ug/g, Mean ± SD) of exDNA and iDNA.

| DNA types | Subtypes | Extracted yields (ug/g) |  |
| --- | --- | --- | --- |
|  |  | Fresh | Pit |
| exDNA |  | 11.91 ± 2.10 | 4.47 ± 0.58 |
|  | Free | 2.66 ± 0.11 | 2.59 ± 0.48 |
|  | Weakly bound | 7.30 ± 2.06 | 0.86 ± 0.15 |
|  | Tightly bound | 1.95 ± 0.36 | 1.01 ± 0.28 |
| iDNA |  | 62.94 ± 6.48 | 28.26 ± 4.15 |

Next, we quantified the abundances of six ARGs (*ermB, sul, tet33, tetG, tetM, tetX*) and three MGEs *(intI1, intI2, intI3)* in all exDNA and iDNA extractions (**Fig. 1**). Significant differences between iDNA and exDNA concentrations in all genes were observed in fresh manure (**Fig. 1A**), with iDNA levels consistently higher than exDNA (p <0.05). Similar trends were observed in pit manure, though only *sul1* and *tetX* genes were significantly more abundant as iDNA relative to exDNA (**Fig. 1B**). To facilitate a direct comparison between solid fresh manure and slurry pit samples, we calculated the exDNA-to-iDNA ratios for each gene (**Fig. 1C**). These ratios were significantly higher in pit than in fresh manure across all genes (Wilcoxon rank sum, p < 0.001), supporting that exDNA is found at higher proportions than iDNA in pit relative to fresh manure samples.

**Figure 1.**
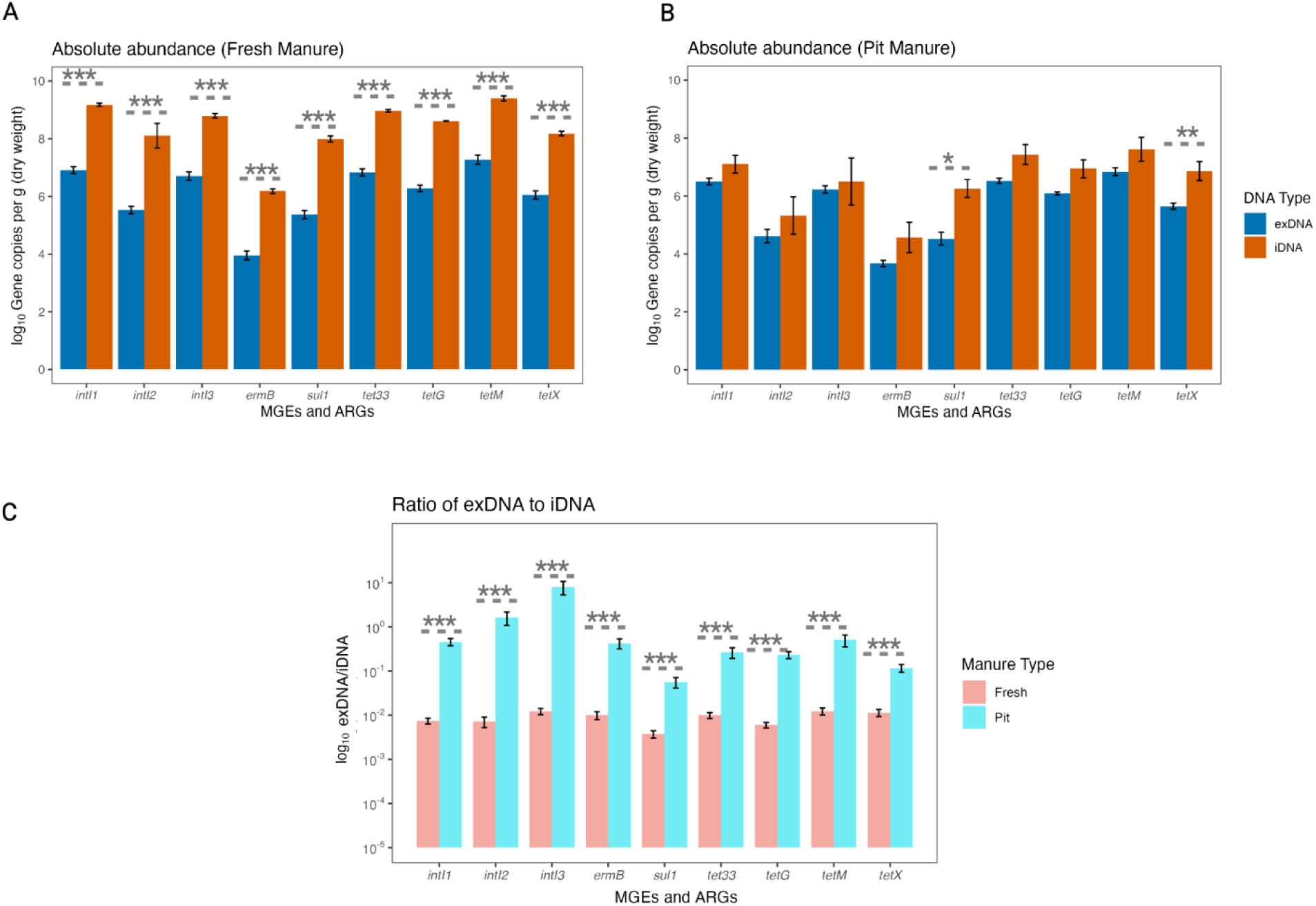
Manure storage pit depletes iDNA genes relative to exDNA (A) Absolute abundance of ARGs’ and MGEs’ copies per g dry weight for Fresh Manure. (B) Absolute abundance of ARGs’ and MGEs’ copies per g dry weight for Pit Manure. (C) Bar plot showing the ratio of exDNA to iDNA in log-scale. Depending on the normality, pairwise significance between the DNA types (exDNA vs. iDNA) or the manure types (Fresh vs. Pit) was determined by independent sample t-test or Wilcoxon sum rank test. Asterisks indicate statistically significant correlations of \**p* < 0.05, \*\**p* < 0.01 and, \*\*\**p* < 0.001, respectively. Error bars represent standard errors of the means (Mean ± SE).

We quantified both free and bound exDNA fractions within the extracted exDNA pool (**Table 1**) and calculated the free-to-bound ratio for fresh and pit samples. For different ARGs, these ratios varied across samples, with no consistent patterns. Significant variations (Wilcoxon rank sum, p <0.01) were found for several genes. Specifically, the ratio was higher in pit manure for *tetM* and, to a lesser extent, *intI2,* while it was higher in fresh manure for *intI3* and *tet33* (**Supplementary Fig. S1**). The *tetM* was the most prevalent (p<0.001) in free form in pit relative to fresh manure.

### Correlation analysis shows strong associations between MGEs and ARGs in DNA categories

To explore the relationship between mobile genetic elements (MGEs) and antibiotic resistance genes (ARGs), we classified detected genes into intracellular (iARGs, iMGEs) and extracellular (exARGs, exMGEs) categories based on their presence in iDNA or exDNA fractions. Correlation network analysis was used to assess co-occurrence patterns among these gene groups. Both fresh and pit exDNA contained 16 strong positive associations (density 0.444, average degree 3.56) but differed in topology: pit-stored manure samples demonstrated less interconnectedness than that of fresh manure, as reflected by a slightly lower clustering coefficient (0.67 vs. 1.00) (**Supplementary Fig. S2,** positive Kendall’s Tau ≥ 0.5).

To increase statistical power and better understand broader patterns of association, data from both fresh and pit samples were combined in subsequent analyses. With the combined data, correlation analysis was used to explore potential relationships between MGEs and ARGs, which may reflect gene mobility or shared selection pressures. Strong associations were observed between iMGEs and iARGs in the iDNA fraction (**Fig. 2A**), particularly among tetracycline resistance genes (*tetX, tetG, tetM*). Notably, *intI1* and *intI3* were strongly correlated with *tetX, tetG, tetM,* and *ermB*, while *intI2* showed a distinct association with *ermB*. In addition, *tet* genes exhibited strong correlations with one another. In the exDNA fraction (**Fig. 2B**), *intI1* and *intI3* were linked to *tetX* and *sul1*, whereas *intI2* exhibited broader associations with *tetX, sul1, tetM,* and *ermB*. Interestingly, *sul1* was exclusively correlated within the exMGEs. Across both intracellular and extracellular environments, only four correlations were consistently significant including *intI1–intI3, intI1–tetX, intI2–ermB, and ermB–sul1*.

**Figure 2.**
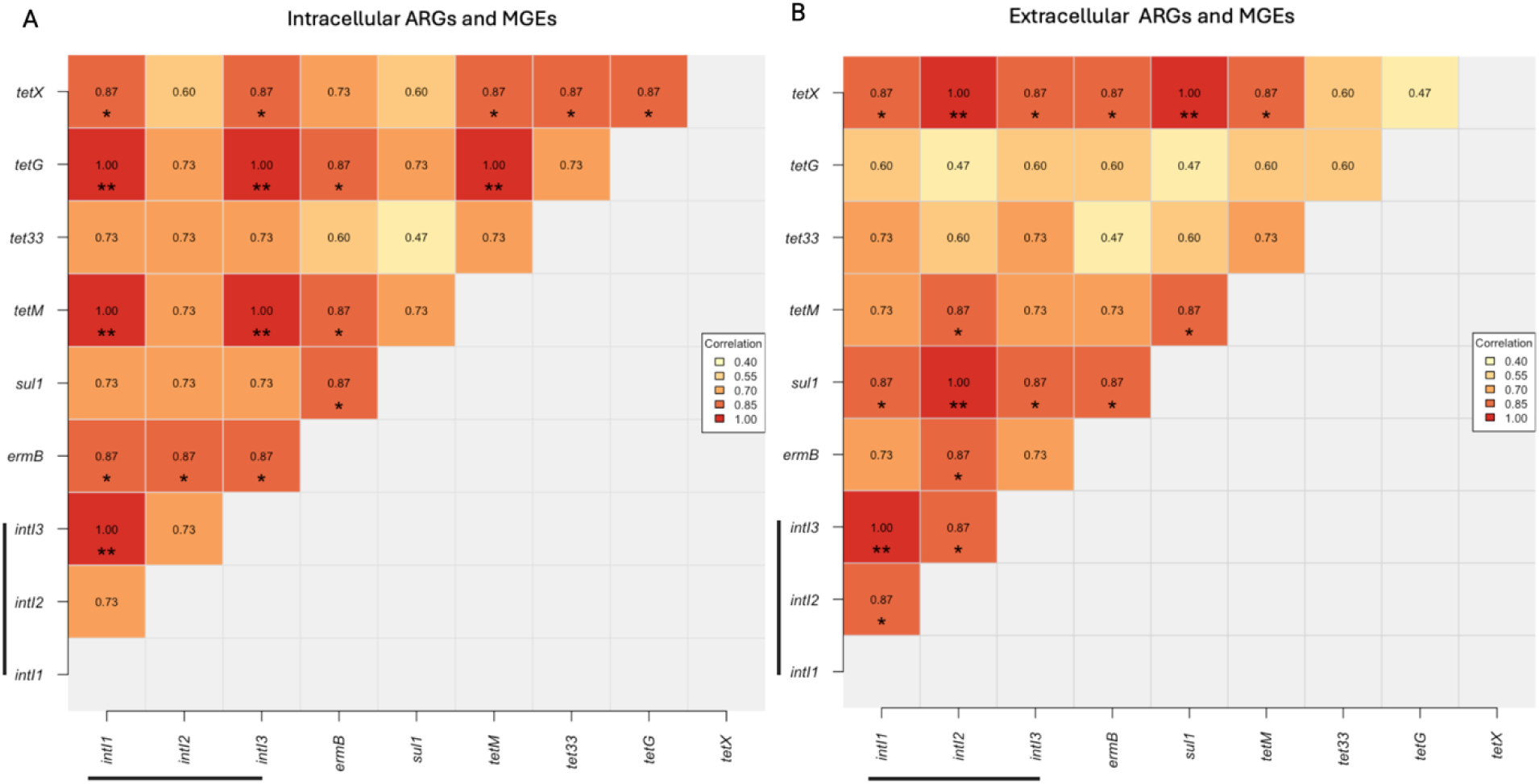
Correlation analysis shows strong associations between MGEs and ARGs in DNA categories. (A) Heat map showing the correlation between iARGs and iMGEs (underlined). (B) Heat map showing the correlation between eARGs and eMGEs (underlined). Values in each square were determined by Kendall’s Tau correlation coefficient. Asterisks indicate statistically significant correlations (* for p < 0.05, ** for p < 0.01).

### Contrasting trends in microbial communities across exDNA and iDNA

To further investigate the community structure in exDNA and iDNA, we performed 16S rRNA amplicon sequencing to characterize the microbial communities across both manure types and DNA categories. Shannon diversity was calculated at the phylum level to assess overall microbial community complexity across sample groups. No significant difference was found between fresh and pit samples (**Supplementary** F**ig. S3A**). Both fresh and pit manure were dominated by Proteobacteria, Firmicutes, and Bacteroidetes. Other phyla such as Actinobacteria, Spirochaetes, and Verrucomicrobia were also detected but in lower abundance (**Supplementary Fig. S3B and S4A).** We also performed Non-metric Multidimensional Scaling (NMDS) analysis (Stress value of 0.06, **Fig. 3**) to assess differences in microbial communities between manure types and DNA fractions. The analysis revealed distinct clustering of iDNA communities from fresh and pit manure, supported by a significant PERMANOVA result (p = 0.001, R² = 63.96%) and indicates strong compositional differences. These differences were further supported at the genus level (**Fig. 3B, see also Supplementary Fig. S4B**); iDNA from fresh manure included genera such as *Ruminobacter*, whereas pit manure was enriched in genera like *Bifidobacterium.* In contrast, exDNA communities from both manure types clustered together, suggesting greater similarity in extracellular microbial profiles across both manure types. The homogeneity of the dispersion was not significantly different between fresh and pit manures (p = 0.75), suggesting that results are likely due to true differences in microbial community composition rather than differences in within-group variability.

**Figure 3.**
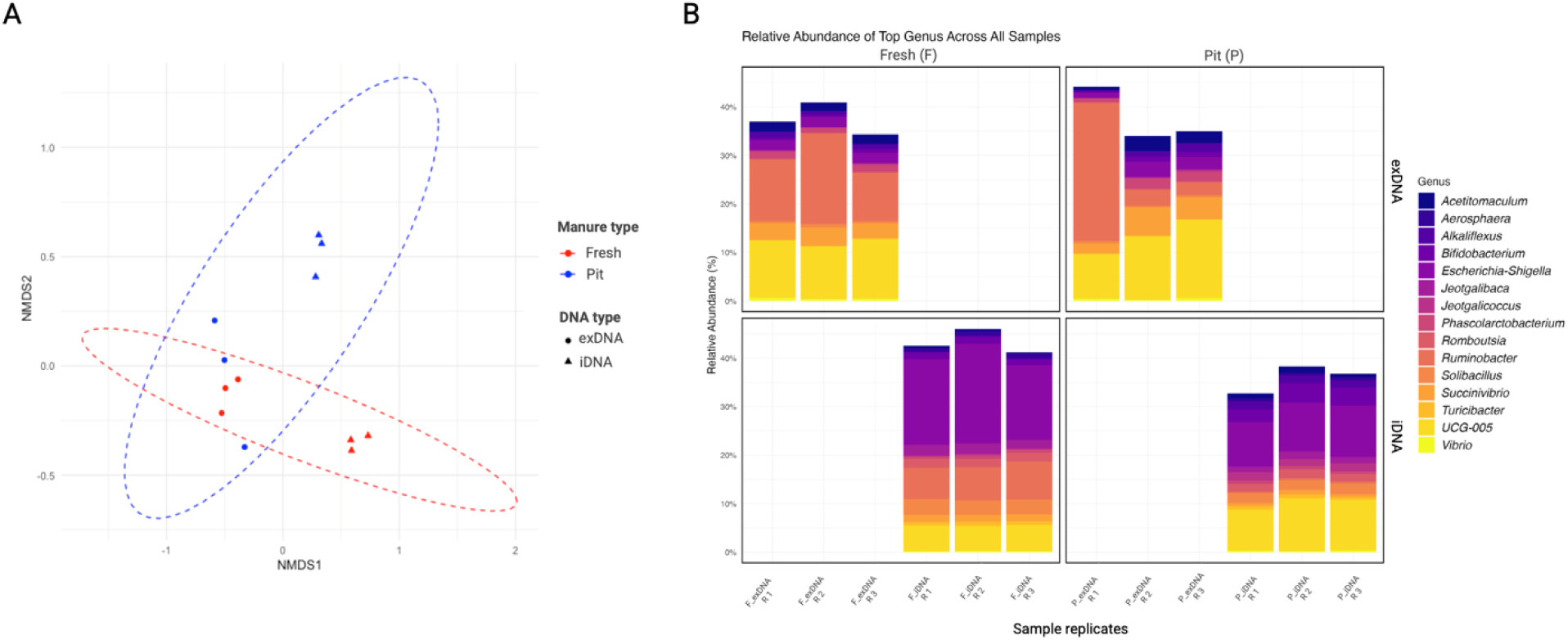
Microbial community analysis shows intermixed exDNA but distinct iDNA communities. A) Non-metric Multidimensional Scaling (NMDS) plot of Bray-Curtis dissimilarity index for taxa abundances between all sample types. Significant clustering was analyzed by PERMANOVA (*p* = 0.001, R² = 63.96%). B) Relative abundance of bacterial genera in exDNA and iDNA. The relative abundance equals the number of reads identified for a specific species by the total number of reads.

## Discussion

This study addresses a critical gap in understanding how exDNA, particularly ARGs within exDNA, change during the storage of dairy manure in pits. Our results indicate exDNA is present in both fresh and pit-stored manures. We find that iDNA consistently exceeded exDNA in abundance across all manure samples, consistent with patterns observed in other nutrient-rich, high-microbial-density environments (Zhang et al., 2013; Sui et al., 2019). However, gene-specific exDNA-to-iDNA ratios revealed relatively higher exDNA levels in pit-stored samples. This enrichment of ARGs within exDNA suggests that manure storage may facilitate the extracellular accumulation of genetic material encoding for antibiotic resistance determinants.

Several factors may contribute to the enrichment of exDNA in manure pits. Short to long-term storage may create conditions that promote microbial cell lysis, releasing intracellular DNA into the extracellular environment. The use of chlorinated disinfectants in dairy wash water can further accelerate this process by inducing oxidative stress and compromising cell membranes, leading to increased DNA release (Fukuzaki, 2006; Du et al., 2015; Guo et al., 2015). Additionally, the anaerobic and nutrient-rich conditions in pits may slow down DNA degradation by limiting nuclease activity, allowing exDNA to persist for extended periods (Nagler et al., 2018b, 2018a). High organic matter and solids like clay minerals, sand particles, and humic substances can also adsorb and protect DNA fragments from enzymatic breakdown (Agnelli et al., 2004; Ceccherini et al., 2009; Pietramellara et al., 2009; Nagler et al., 2018a). Our observations that exDNA may be enriched in manure pits combined with these environmental conditions highlight this environment as an area as a reservoir for exARGs.

The physicochemical properties of dairy manure likely govern the retention, protection, and mobilization of ARGs via exDNA, as reflected by the varying free-to-bound ratios observed in the samples. Pit samples showed greater ratios of free-to-bound exDNA specifically for *tetM*. This pattern suggests that *tetM* may be more prevalent in a free and potentially more accessible form compared to its presence in fresh manure samples. This availability in dairy pits could potentially increase its association with horizontal gene transfer. Conversely, genes such as *intI2*, *intI3*, and *tet33* were more bound in pit samples than in fresh manure. Together, these observations suggest that ARGs may vary in their potential to be in extra- or intracellular DNA pools. The degree to which an ARG is bound in DNA pools can also affect the persistence of these determinants in the environment as manure is applied to agricultural soils. Previous studies have shown that exDNA can persist in soils for extended periods and that transformation can occur rapidly under favorable conditions, even with relatively low exDNA concentrations (Levy-Booth et al., 2007; Pietramellara et al., 2009; Kittredge et al., 2022; Xin et al., 2025), and thus our results present a strong rationale for further study on the enrichment of exDNA in manure pits and its persistence both in the pit and downstream in land application.

Our observation that integrons were enriched in exDNA in pit manures is significant because these genes are often embedded within mobile elements (Yang et al., 2020; Buta-Hubeny et al., 2022). While they typically reside on chromosomes, environmental pressures from pollutants like antibiotics and heavy metals can force their mobilization, especially on plasmids (Gillings et al., 2015). In our study, *intI1* was found to have the highest absolute abundance among integron genes. This observation aligns with previous findings reporting *intI1* in as abundant in both exDNA and iDNA copies in various anthropogenic and agricultural waste sources (Guo et al., 2018; Dong et al., 2019). The persistence of extracellular *intI1* has also been abundant in samples from wastewater treatment plants (Wang et al., 2020). Moreover, our study found *sul1* associated with integrons in exDNA. This finding is significant given that *sul1* is often part of the 3’-conserved segment of class 1 integrons (Jiang et al., 2019; Lima et al., 2020). Its presence in exDNA suggests a potential for horizontal gene transfer via plasmids and is consistent with previous studies showing that *sul1*-containing plasmids (e.g., IncF plasmid) can be conjugally transferred (Jiang et al., 2019). Recent research also identified an increase in *sul1* in the exDNA pool across wastewater treatment plant systems (Martínez-Quintela et al., 2024). In iDNA pools, we observed a correlation between *intI2* and *ermB*. This association implies that *ermB*, which confers resistance to macrolide antibiotics, may be co-selected with integrons. The association of *ermB* with integrons (i.e., *intI1*) has also been previously reported in freshwater iDNA samples (Sabatino et al., 2025). The presence of this co-located arrangement in iDNA and not exDNA suggests that the genetic linkage may be maintained within living bacteria. The presence of this co-located arrangement in iDNA and not exDNA suggests that the genetic linkage is actively maintained within living bacteria (Partridge et al., 2018).

Understanding the microbial community structure in exDNA and iDNA pools provides an additional context for interpreting ARG dynamics and their potential for horizontal transfer. Although 16S rRNA is typically associated with intact bacterial cells, its detection in exDNA is common and may result from recent cell lysis, active secretion as part of biofilm formation, or other physiological processes such as outer membrane vesicle release (Guo et al., 2018; Dong et al., 2019; Martínez-Quintela et al., 2024). Our results indicate distinct beta diversity patterns across DNA pools from fresh and pit manure sources. While both fractions share similar taxa, their overall community structures differ substantially. This result likely reflects differences in selective pressures, which are consistently acting on iDNA from living cells, but not on exDNA, which comes from lysed cells and is less affected by the surrounding environmental forces (Nagler et al., 2021, 2022). The differences in iDNA between fresh and pit manure reflect their distinct environments. In our study, we detected enriched *Ruminobacter* in fresh and *Bifidobacterium* in pit iDNA. These results support that fresh manure originates from the gut of the animal and is fiber-rich, and supporting genera like *Ruminobacter* that thrive on rumen fermentation. In contrast, the pit manure goes through fermentation and storage, creating conditions that encourage the growth of more adaptive genera like *Bifidobacterium* (McGovern et al., 2020; Sukhum et al., 2021). Conversely, the exDNA pools from both manure types tend to be more similar, likely because exDNA binds to organic matter, which protects it and preserves a kind of historical snapshot of the microbial community and may be less affected by ongoing selective pressures (Nagler et al., 2021).

Although our findings provide important initial insights into exDNA pools in dairy manures, several limitations should be acknowledged. We could not completely eliminate iDNA contamination from our exDNA; however, minimal false positives (gfp recovery of 0.003%– 0.02%) were lower than the 1.3% reported previously (McKinney and Dungan, 2020). The presence of the *gfp* gene in exDNA fractions from whole cells of *E.coli* containing the plasmid is likely due to the excreted DNA from live or partially lysed cells. Other studies that used membrane filters to remove microbial cell contamination from exDNA primarily focused on water samples (Corinaldesi et al., 2005; Alawi et al., 2014; Zhang et al., 2018). In contrast, our study used manure samples, which could trap exDNA on the filters during the filtration step, so this method was avoided. We also acknowledge that the limited sample size in this study was a constraint, and future research should include a larger number of samples collected across multiple farms, locations, and timepoints to improve representativeness.

## Conclusion

This study provides new insights into the dynamics of exDNA and its ARGs during dairy manure storage. We demonstrate that manure pits can act as reservoirs for exDNA and suggest that storage conditions, including physicochemical properties and microbial processes, may influence the persistence and potential mobility of ARGs. The enrichment of exDNA in pits raises important questions about its role in horizontal gene transfer in both the pit and manure land application, particularly given the presence of mobile genetic elements such as integrons and plasmids and their association with exARGs. Addressing these knowledge gaps through future studies is essential to deepen our understanding of exDNA dynamics and ARG mobility under varying storage conditions and management practices.

## Acknowledgements

This project was supported by AFRI food safety grant no. 2021-68015-33495 from the USDA National Institute of Food and Agriculture.

## Data availability

All data are available in GitHub through Zenodo (https://doi.org/10.5281/zenodo.17544973).

## Authors’ contribution

Study planning: NS and AH; Study plan validation: All authors; Data collection: NS and DA; Data analysis: NS; Data interpretation: NS, AH, and LJ; First drafting: NS; Review and re-drafting: NS. AH and LJ; Final approval: All authors.

---

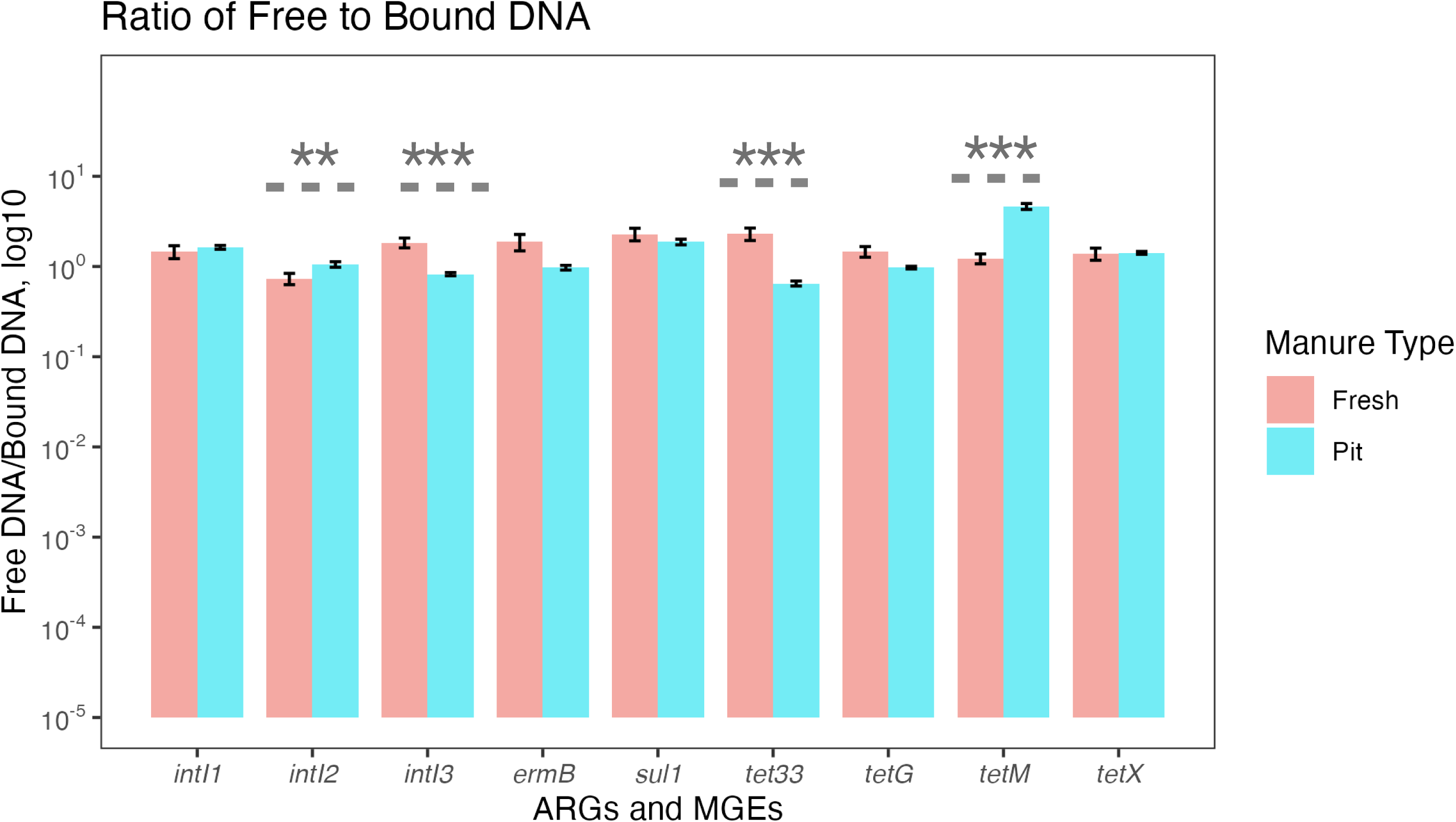

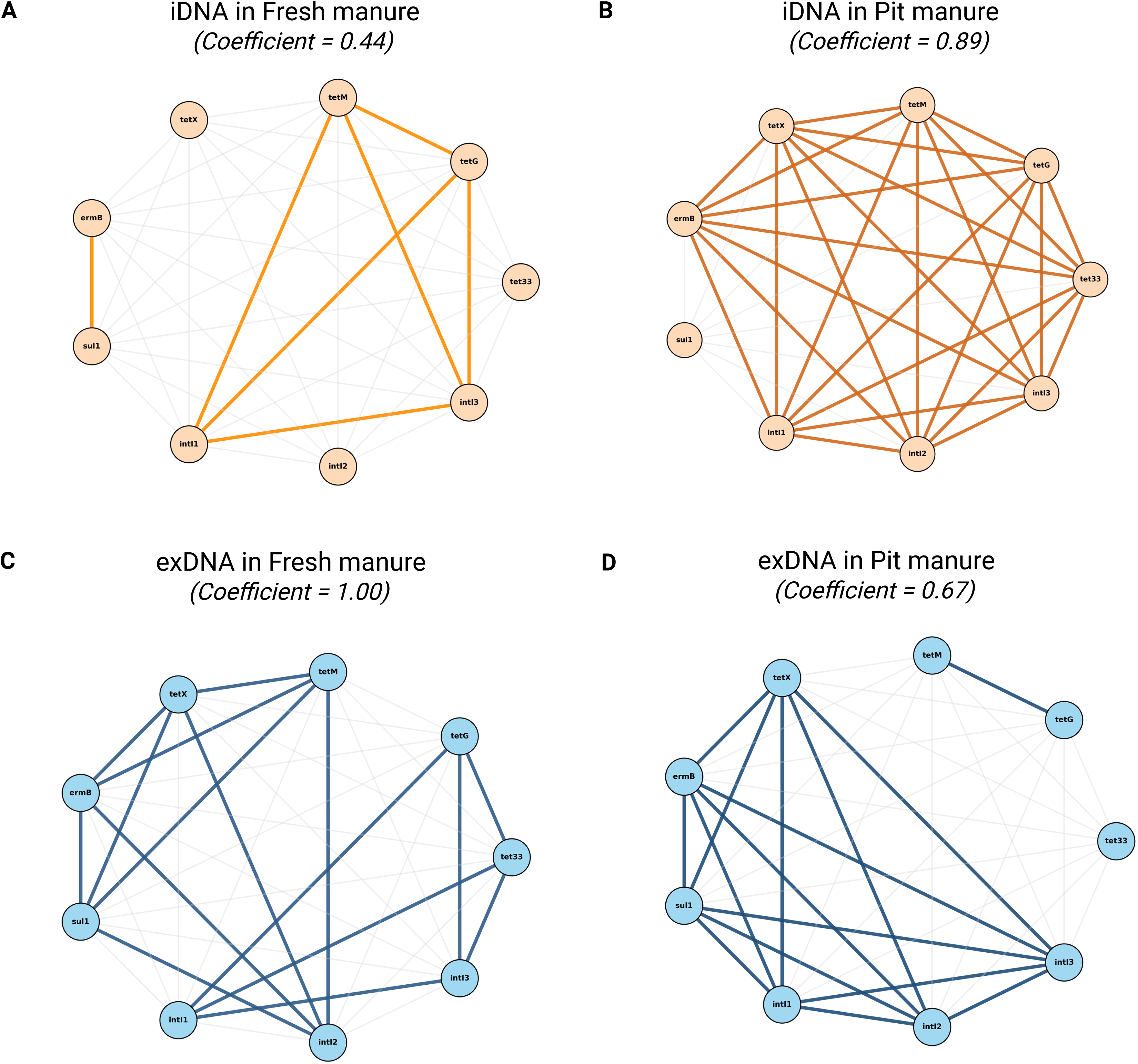

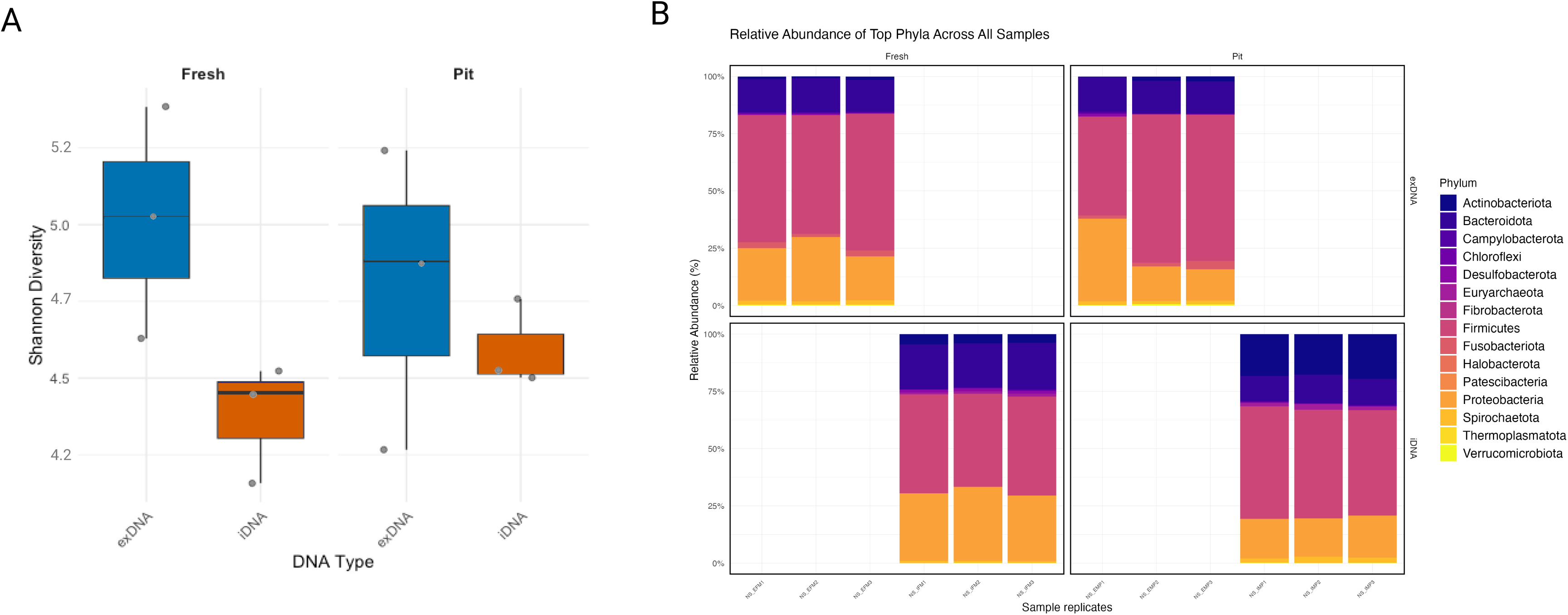

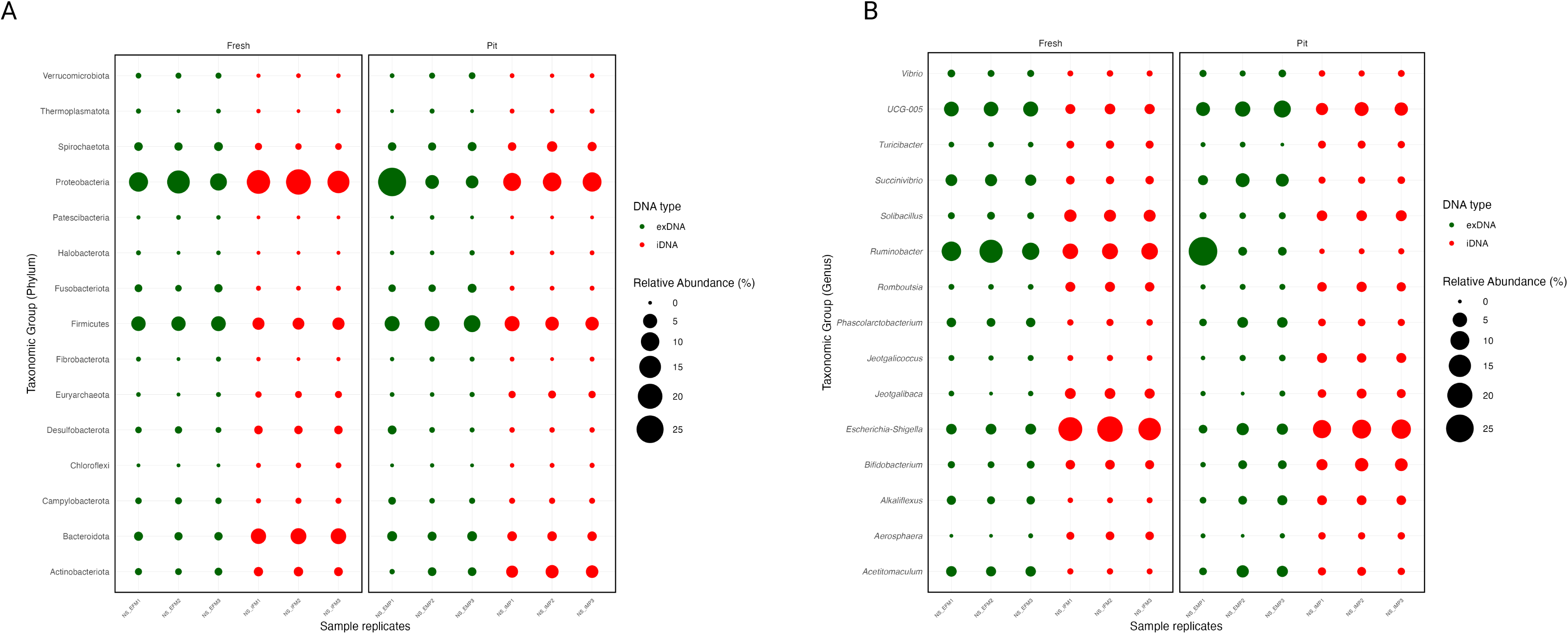

